# CRISPR-mediated knockdown of oxytocin receptor in extended amygdala reduces stress-induced social avoidance in female California mice

**DOI:** 10.1101/2025.06.10.658921

**Authors:** Valentina Cea Salazar, Arjen J. Boender, Adele M. H. Seelke, Liam Gaard, Sabrina L. Mederos, Sophia Rogers, Xiomara Z. Gutierrez, Karen L. Bales, Larry J. Young, Brian C. Trainor

## Abstract

Oxytocin receptors (OTRs) within the extended amygdala and nucleus accumbens (NAc) have been implicated in modulating social behaviors, particularly following stress. The effects of OTR could be mediated by modulating the activity of pre-synaptic axon terminals or via receptors in post-synaptic neurons or glia. Using a viral-mediated CRISPR/Cas9 gene editing system in female California mice (*Peromyscus californicus*), we selectively knocked down OTR in the anteromedial bed nucleus of the stria terminalis (BNST) or NAc to examine their roles modulating social approach and vigilance behaviors. Knockdown of OTR in the BNST attenuated stress-induced decreases of social approach and had less robust effects on vigilance when interacting with a target mouse behind a wire barrier. In this large arena, where mice could control their proximity to a target mouse, BNST OTR knockdown also increased investigation of a non-social stimulus (empty cage). Behavioral effects of BNST OTR knockdown were weaker in the small arena where focal mice physically interacted with target mice. Interestingly, OTR knockdown in the NAc, reduced stress-induced social vigilance without affecting social approach. These effects could mediated altered encoding of socially aversive experiences, as knockdown manipulations were performed before stress exposure. Together, these results highlight effects of local OTR on social behavior are region-specific.

## INTRODUCTION

Individuals affected by anxiety disorders experience significant distress in social situations that can lead to self-isolation and avoidance behaviors (Abramowitz & Deacon, 2010). Women are twice as likely to develop anxiety disorders compared to men (Kessler et al., 2012). While there are existing pharmaceutical and behavioral therapeutic interventions, ∼40% do not respond to these treatments (Williams & Trainor, 2018). Oxytocin is a neuropeptide that modulates social and emotional behaviors (Gimpl & Fahrenholz, 2001) and has been considered as a potential alternative therapeutic to existing antidepressants (MacDonald & Feifel, 2013 and 2014; Modi & Young., 2012; Gutkowska & Jankowski, 2012; Giovanna et al., 2020; van Zuiden et al., 2017; Schultebraucks et al., 2022; Rashidi et al., 2025). However, the field has struggled to explain findings where intranasal oxytocin sometimes decreases anxiety symptoms but in other cases increases anxiety and other psychiatric symptoms (Tabak et al., 2022; Schultebraucks et al., 2022). A potential explanation for these results is the social salience hypothesis of oxytocin (Bartz et al., 2011; Shamay-Tsoory & Abu-Akel, 2016), which proposes that oxytocin can modulate behaviors by enhancing attention to both positive and negative social contexts. Several lines of evidence indicate that oxytocin promotes salience by acting in discrete neural circuits to drive these dynamic responses (Steinman et al., 2016 and 2019; Nasanbuyan et al., 2018; Osakada et al. 2024; Duque-Wilckens et al., 2018).

In appetitive social contexts, activation of oxytocin receptors (OTR) in the mesolimbic dopamine system plays a key role in promoting social approach behaviors. The nucleus accumbens (NAc) modulates motivational and learning processes (Salgado & Kaplitt, 2015), and activation of OTR within the NAc facilitates social approach (Williams et al., 2020), social learning (Nardou et al., 2019), pair bonding (Liu & Wang, 2003; Rigney et al., 2025) and social novelty seeking (Smith et al., 2017). Activation of OTR in the ventral tegmental area (VTA) has also been found to modulate social motivation (Borland et al., 2017) and learning (Hung et al., 2017). It was hypothesized that oxytocin acting in the mesolimbic dopamine system facilitates social salience in aversive social contexts (Shamay-Tsoory & Abu-Akel, 2016). However, recent evidence suggests that OTR in the extended amygdala may play an important role during stressful social contexts. Social defeat stress activates a population of oxytocin neurons in the medioventral bed nucleus of the stria terminalis (BNST) in *Peromyscus californicus* (California mice) (Steinman et al., 2016) and C57Bl/6J mice (Nasanbuyan et al., 2018). The BNST modulates anxiety-related and social behaviors (Lebow & Chen, 2016) and several subregions have strong molecular (Gegenhuber et al., 2022) and anatomical (Campi et al., 2013; Allen & Gorski, 1990) sex differences. Antisense knockdown of oxytocin in the BNST prevented defeat stress-induced social avoidance and social vigilance in female California mice (Duque-Wilckens et al., 2020). These BNST oxytocin neurons project to the anteromedial BNST (BNSTam) where activation of OTR promotes avoidance and vigilance (Duque-Wilckens et al., 2018; Luo et al., 2022). Oxytocin can also promote aversive responses in non-social contexts, as oxytocin in the dorsolateral BNST (BNSTdl) enhanced fear-potentiated startle (FPS) response in rats (Moaddab & Dabrowska, 2017). Together these data suggested that distinct neural circuits mediate the effects of oxytocin in appetitive and aversive social contexts (Steinman et al., 2019).

Most of the studies reviewed above relied on pharmacological manipulations to target OTR function. Pharmacological approaches target both locally expressed post-synaptic receptors and pre-synaptic receptors. Importantly, at least some behavioral effects of OTR can be mediated by pre-synaptic receptors. Dölen and colleagues showed that the behavioral effects of OTR antagonists infused into the NAc were mediated by OTR expressed on presynaptic terminals originating from the dorsal raphe nucleus (Dölen et al., 2013; Nardou et al., 2023). These studies were performed in C57Bl/6J mice and utilized transgenic mouse lines to specifically target different populations of OTR. The CRISPR/Cas9 gene-editing systems provides a mechanism to perform similar manipulations in species for which transgenic lines are not available. A viral-mediated CRISPR/Cas9-based tool was developed to target OTR expression across a variety of rodent species used as models to study social behavior (Boender et al., 2023). When a guide RNA targeting the *Oxtr* gene was expressed with Cas9, OTR binding was reduced across six different rodent species, including California mice. We used this gene editing system to determine whether locally expressed OTR in the BNST or NAc modulate social approach and social vigilance in female California mice. This viral construct was recently used in a male spiny mouse model in the NAc, which reduced huddling behaviors with strangers as well as some feeding behaviors identifying a role for OTRs in non-reproductive contexts (Fricker et al., 2025).

California mice are monogamous rodents in which paired males and females defend joint territories. The high levels of aggression in female California mice make this species ideal for studying the effects of social defeat stress (SDS) in both males and females without the use of CD1 aggressive male mice generally used in C57BI/6J models (Steinman & Trainor, 2017; Lake & Trainor, 2024). In this study we assessed the impact of OTR knockdown on behavior before and after stress using two behavioral assays in female California mice. Focal mice were tested in a large arena with an unfamiliar female stress-naïve target mouse confined to a small wire cage, which allows for precise measurements of social approach and social vigilance. Social vigilance is a risk assessment behavior in which an individual avoids but attends to a social context (Walker et al., 2023; Wright et al., 2020). Social approach is assessed by time spent within one body length of the cage. Focal mice were also tested in a smaller arena with freely moving stress-naïve and aggressive SDS target mice. This test is ideal for the observation of affiliative and defensive behaviors. Previous studies showed that social approach and social vigilance are modulated by OTR acting within complementary networks that include the BNST and NAc (Duque-Wilckens et al., 2018 and 2020; Luo et al., 2022; Williams et al., 2020). We used a CRISPR-mediated virus to mutate the coding region of the *Oxtr* gene to induce a reduction in OTR receptor binding in female California mice. We refer to reductions of OTR binding in CRISPR treated groups as “knockdown” within a specific brain region and examined whether changes in locally expressed OTR corresponded to changes in social behavior before and after social defeat stress. We used two behavioral assays to assess the roles of OTR on social approach as well as ethological measures of social interaction.

## MATERIALS AND METHODS

### Animals

All studies were conducted using California mice at the University of California, Davis (UCD). Protocols were approved by the UCD Institutional Animal Care and Use Committee (IACUC). Adult (90 days old) female California mice were studied. Mice were housed with same-sex cagemates after weaning in groups of 2 to 4 within clear polypropylene cages. Cages were set up with Sani-Chip bedding (Harlan Laboratories, Indianapolis, IN, USA) and 2 nestlets per cage (Ancare, Bellmore, NY, USA). Mice were kept on a 16L:8D light cycle and had *ad libitum* access to food and water. Viral infusion/surgery were conducted in light cycle, while behavioral testing and stress exposure were performed during the dark cycle.

### Intracranial surgeries

Anesthesia was induced via exposure to a 5% isoflurane mixture with oxygen and then maintained at 2-3.5%. With a stereotaxic apparatus, mice were bilaterally injected 300nL with either the *Oxtr* guide RNA virus or a scramble sequence virus. Mice were randomly assigned to treatment group (CTRL/KD). Coordinates were calculated using the online California mouse brain atlas at brainmaps.org. For NAc we used, AP: +0.51, ML: ±1.5, DV: -6.0. For BNST we used, AP: + 0.29, ML: ± 1.1, DV: -5.85. Mice assigned to OTR knockdown received viral cocktail consisting of an AAV9 vector expressing *Streptococcus pyogenes* Cas9 under the RSV promoter (Fig. 1A, AAV9-RSV-spCas9,stock titer 1.5 x10^10^ genomic copies/ul; injected titer 7.5 x 10^9^ gc/ul) and a second AAV9 vector encoding a guide RNA targeting Oxtr (AAV9-U6-gRNA(ΔOXTR.1)-CMV-eGFP, 5′-GGTGCTTCATGAAAAAGAAG-3′, stock titer 2.0 x 10^10^gc/ul stock; injected titer 1.0 x 10^10^ gc/ul) mixed in a 1:1 (Boender et al., 2023).This guide RNA targets a sequence of the *Oxtr* gene that is evolutionarily conserved across several rodent species. Control mice received a cocktail consisting of the Cas9 vector and a scrambled gRNA sequence. All mice are given a carprofen injection at the beginning of surgery (1.45 mg/kg dosage) and then 3 daily doses during recovery. Skin staples were used to close the skin and were removed 7 days post-surgery.

**Figure 1:**
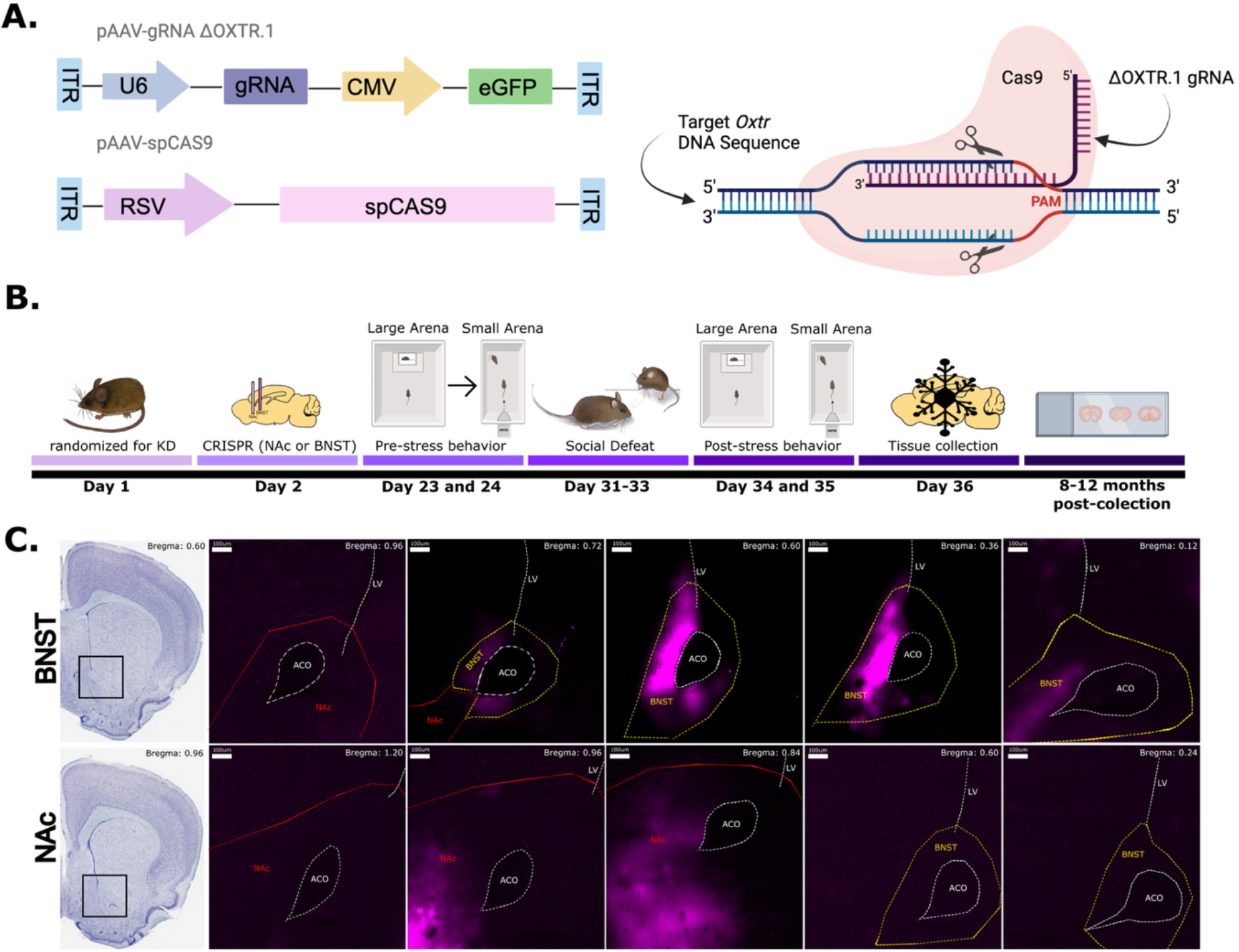
The CRISPR/Cas9 genome-editing system can selectively target the BNST and NAc in female California mice. (A) Schematic of AAV plasmids and CRISPR mechanism of action. (B) Experimental timeline. (C) Representative photomicrographs of viral spread in the BNST and NAc knockdown groups. Scale bar: 100um. LV, lateral ventricle; ACO, anterior commissure; NAc, nucleus accumbens; BNST, bed nucleus of the stria terminalis.

### Behavioral timeline

Three weeks after surgery, all mice were tested in a Large Arena Social Interaction test (Fig. 1B). In prairie voles 2 weeks of viral expression are sufficient to decrease OTR binding (Boender et al. 2023), so we are confident that mice assigned to knockdown had reduced OTR expression when these behavior tests were conducted. One day later mice were tested in a small arena social interaction test. In this test target mice are not caged, allowing direct contact with focal mice. Three days after these behavioral tests all focal mice underwent three days of social defeat on consecutive days. One week later both the large and small arena social interaction tests were repeated as described above. Mice were never tested with the same Target or aggressive Target mouse (except for post-stress interaction with aggressive Target) in the large or small arena tests.

### Large arena test

This test consisted of 3 phases: Open Field, Acclimation, and Interaction (Greenberg et al. 2014). Focal mice were placed in the large rectangular arena, made opaque white acrylic (89 × 63 × 60 cm). Each phase lasted 3 min and was recorded via AnyMaze software. In the Open Field phase, mice were allowed to explore the empty arena. Time spent in the center of the arena and total distance traveled was recorded. This was followed by the Acclimation phase where an empty wire cage (14×17×14.5 cm) was placed on one end of the arena. Finally, in the last phase an unfamiliar stress-naïve female target mouse was placed inside the wire cage. This test was repeated 1 week after social defeat stress. A different target mouse was used from the pre-stress test to post-stress testing.

### Small Arena test

The small arena test was conducted in an acrylic chamber (51 × 25.4 × 76 cm) connected to a small chamber with a sliding door for introducing target mice. In the Open Field phase, focal mice entered through the attached chamber (13 × 10 × 18 cm) and explored the empty arena for 3 minutes. We then assessed interactions with target mice with (aggressive) or without (naïve) prior experience winning aggressive encounters (Wright et al., 2023; Luo et al., 2024). First a female naïve target mouse was introduced to the arena and interactions with the focal mouse were recorded for 3 minutes. The target mouse was removed and then an aggressive female target mouse was introduced to the arena for final 3 minutes. This test was repeated 1 week after social defeat stress (Fig. 1B). In the post-stress tests, different target mice were used from the pre-stress tests. However, each focal mouse had interacted with the aggressive target mouse from social defeat encounters. All videos were scored using BORIS by a single trained observer without knowledge of the treatment group status of the focal mouse. Here affiliative (nose-to-nose sniffing, anogenital sniffing, and bodily sniffing), coping (auto-grooming and flipping), defensive (lunging), and behaviors were recorded. Social approach was not assessed in this test due to the size of the arena and the lack of machine learning to score “passing” behavior versus “approach.” Flipping behavior is a motivated behavior in California mouse that has been identified as a coping behavior (Minie et al., 2021).

### Social defeat stress

After testing in the large and small arena tests, all focal mice underwent social defeat stress. Each mouse experienced 3 consecutive days of social defeat stress as previously described (Greenberg et al., 2014). On each day, the focal mouse was placed into the home cage of a sexually experienced aggressive resident same-sex mouse. Aggressive females are paired with a vasectomized male as this results in more consistent territorial aggression (Trainor et al. 2013). Each episode of this defeat lasts 7 minutes or until the intruder/experimental mouse is attacked/defeated a total of 7 times, whichever comes first. After testing, experimental mice are immediately placed back in their homecage. Post-stress testing was conducted one week after the last episode of defeat. This 3-day protocol does not induce physical injury to focal or resident mice and results in behavioral changes that can be observed up to 10 weeks after the last episode of defeat (Steinman et al., 2016; Trainor et al. 2011).

### Histology and Autoradiography

After post-stress behavior testing mice were anesthetized with isoflurane, decapitated and brains were removed to be flash frozen on dry ice. Brains were stored at -80° C and then cut with a cryostat at 20μm and mounted directly on frost plus slides in series. One set of slides was used to determine viral spread by assessing green fluorescent protein (GFP) expression (Fig 1C). These images were used to determine spread of the virus. Adjacent sections were saved on an alternate of slides for OTR autoradiography (as in Hartman et al., 2018). Briefly, slides were removed from -80°C storage, thawed at room temperature for 1 h, fixed in a light 0.1% paraformaldehyde (pH 7.4) solution, and washed twice in Tris Buffer, before being incubated in the tracer buffer at room temperature for 60 min. Tracer buffer consisted of 50 mM

Tris-HCl buffer (7.4 pH) with 10 mM MgCl_2_, 0.1% bovine serum albumin, and 50 pM of radiotracer. For OTR binding, [125I]-ornithine vasotocin analog [^125^I-OVTA] [vasotocin, d(CH_2_)5[Tyr(Me)2, Thr4, Orn8, (125I)Tyr9-NH2]; 2200 Ci/mmol] was used (Revvity, Waltham, MA, USA). Previously we observed that the CRISPR tools we used in this study reduced OTR binding but not V1aR binding in California mice (Boender et al., 2023). Following the incubation period, unbound radioligand was removed by 4 washes in 50 mM Tris buffer and 2% MgCl_2_, pH 7.4, and then dipped in dH_2_0 and air dried.

The slides were dried overnight and exposed to Cytiva Amersha, Hyperfilm (Cytiva Life Sciences, Washington, DC, USA) for 4 days with a set of ^125^I standard microscales (American Radiolabeled Chemicals, St. Louis, MO, USA). We then used MCID Core Digital Densitometry system (Cambridge, UK) to quantify *Oxtr* binding in each target brain. For each specimen, optical binding density (OBD) values were calculated for each region of interest (ROI), as well as one background area where no binding was detected. Three separate measurements for each ROI were taken per specimen and averaged. For each specimen, the average background binding value was subtracted from each average ROI measurement to yield normalized OBDs across specimens. There were 2 samples that were damaged in the slide processing and treatment designation for knockdown was determined with GFP photomicrographs of coronal sections of NAc and BNST. Based on oxytocin receptor binding and GFP spread, two mice were reassigned treatment groups (one NAc to BNST, one BNST to NAc) and four mice had OTR binding levels in the BNST and NAc that were below the lowest value observed in control groups. Based on these observations these mice were reassigned to a NAc + BNST knockdown group. We limit our inferences from this group due to the small sample size.

### Statistical analysis

Autoradiography data were analyzed using one-way ANOVA in R with planned comparisons to compare treatment groups with controls. For large arena tests behavior was analyzed with one-way ANOVA followed by planned comparisons. We used paired t-test within each treatment group to test if there were differences before versus after stress. For the small arena tests each behavior, we conducted separate analyses by social partner type (naive vs. aggressive target) and timepoint (pre-stress vs. post-stress). For small arena tests each dependent variable was tested across treatment groups using nonparametric Kruskal-Wallis test with pairwise comparisons using the Mann-Whitney U test.

## RESULTS

### OTR Knockdown in the BNSTam and NAc in Female California Mice

Autoradiography analysis (Fig. 2A) showed that *Oxtr* CRISPR altered OTR binding in the BNSTam (Fig. 2B, F_3,26_ = 13.69, p<0.001) and NAc (Fig. 2C, F_3,26_= 12.18, p<0.001). Mice assigned to BNST knockdown had reduced OTR binding in BNST (Fig. 2B, planned comparison, P<0.001, Cohen’s d =2.36) but not NAc (Fig. 2C) compared to controls. In contrast, mice assigned to NAc knockdown had reduced binding in NAc (Fig. 2C, planned comparison p<0.001, d=0.48) but not BNST (Fig 2B) compared to controls. In some animals binding was reduced in both the BNST and NAc, so these animals were included as a separate group as “BNST + NAc KD.” There were no differences in OTR binding in the LS (Fig. 2D), suggesting that our manipulations were site-specific.

**Figure 2:**
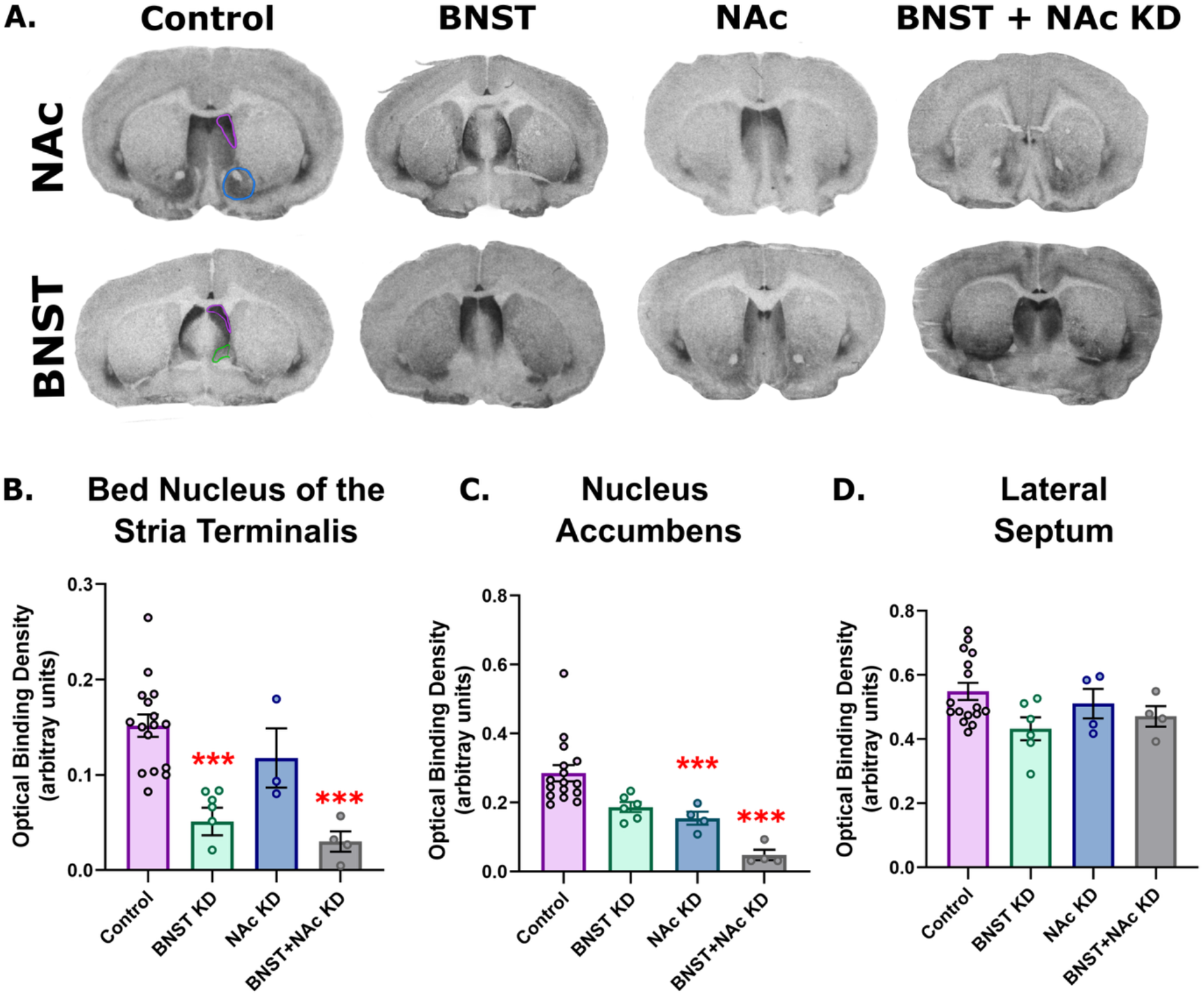
Inhibition of *Oxtr* expression in female *Peromyscus californicus* in the extended amygdala and nucleus accumbens. (A) Representative images of *Oxtr* manipulation in the BNST and NAc with OTR autoradiography. Control panels show regions quantified for NAc (blue), BNST (green) and LS (purple). (B) Control female mice compared to CRISPR-based knockdown of BNST and other treatment groups. Controls have significantly higher OBD than the BNST KD group (P= <0.001). (C) Control female mice compared to CRISPR-based knockdown of NAc and other treatment groups. Controls are significantly higher from the NAc KD group (P<0.001). (D) Controls compared to all groups for density comparison in “control” region, the lateral septum. *** P < 0.0001 control vs. knockdown.

### OTR Knockdown within the BNSTam Prevents Stress-Induced Changes in Social Approach and Vigilance in the Large Arena

Three weeks after bilateral CRISPR injections, adult female California mice were tested in the large arena test to measure baseline behaviors before and after social defeat stress (Fig. 3A). In pre-stress observations there were no differences in social approach (Fig. 3B) or vigilance (Fig. 3C). After stress, OTR knockdown showed a trend to affect social approach behavior (F_3,27_ = 2.50, p = 0.07). Planned comparisons showed that BNST (p=0.04, d=0.43) and BNST+NAc (p=0.04, d=0.01) knockdown groups had higher social approach compared to controls (Fig. 3B). For social vigilance there were significant differences between groups after stress (Fig. 3C, F_3,27_ = 3.50, p = 0.03). Planned comparisons showed lower vigilance in the NAc group (p=0.01, d=0.26) and a trend for lower vigilance in the BNST+NAc knockdown (p=0.052, d=0.30) compared to controls. Decreased vigilance in the BNST knockdown group compared to control was also at trend level (p=0.06, d=0.44). In paired comparisons, controls before and after stress increased their vigilance (p=0.001, d=-1.19) and decreased their approach (p=0.008, d=0.82). Effects of stress were not significant in the other treatment groups. To determine whether the extent of OTR knockdown affected behavior after stress, we examined correlations between receptor density and social approach in the BNST knockdown group and controls (Fig. 4). Receptor density was not significantly correlated with social approach (Fig. 4A, Control: R=-0.16, p=0.56; BNST KD: R=-0.17, p=0.71) or vigilance (Fig. 4B, Control R=-0.001, p=0.98; BNST KD R=0.121, p=0.8). In contrast, a strong negative correlation between social approach and vigilance was observed after stress in controls (R=-0.640, p=0.007), whereas the same relationship in BNST KD mice showed a similar relationship that did not reach significance (R=-0.682, p=0.09).

**Figure 3:**
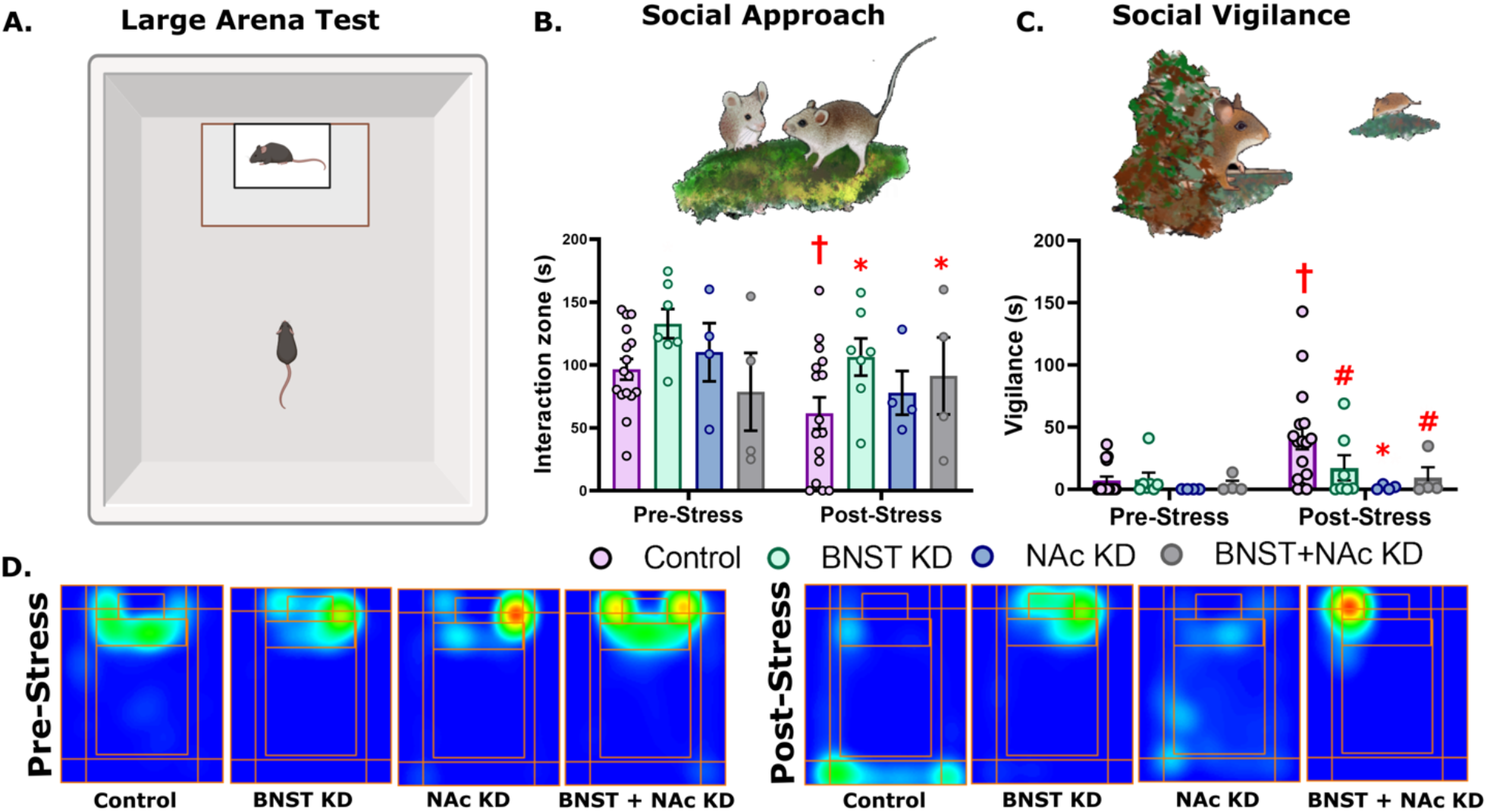
Oxytocin receptor knockdown increases social approach and decreases vigilance after stress in the large arena test. (A) Schematic illustration of the Large Arena Social Interaction test. (B) Effects of OTR knockdown on social approach before and after social defeat stress (BNST KD n= 7, NAc KD n= 4, BNST + NAc KD, n= 4, Control n= 16). (C) Effects of OTR knockdown on social vigilance before and after social defeat stress (D) Representative heatmaps for the interaction phase showing reduced social approach determined by time spent in the interaction zone in controls but not other treatment groups.* P < 0.05 vs. control, # p=0.06 for BNST KD vs. control, # p=0.052 for BNST +NAc KD vs. control, † vs. pre-stress paired comparison. Drawings by Dr. Natalia Duque-Wilckens.

**Figure 4:**
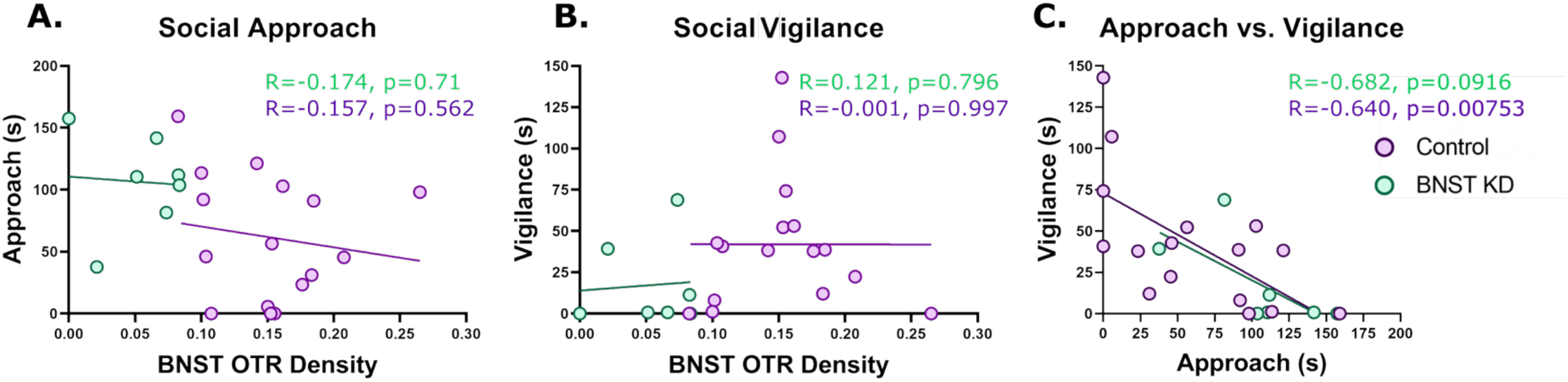
Correlational analysis of social approach and vigilance for controls and BNST KD. (A) OTR BNST density (OBD) compared to social approach behavior after stress. (B) OTR BNST density (OBD) compared to social vigilance behavior after stress. (C) Social approach compared to social vigilance after stress.

There were no differences in time spent in the center of the arena or total distance in the open field phase (Fig. 5 B and C). During the acclimation phase, when the target mouse was absent, there were no differences in approach to the empty cage before stress (Fig. 5D). After stress, approach to the empty cage in control mice did not differ from before stress levels, suggesting that mice did not habituate to the cage. There were effects of knockdown after stress (F_3,27_=3.920, p=0.02). Mice with BNST knockdown showed higher levels of approach to the empty cage compared to controls (p=0.01, d=-0.42), whereas NAc KD and BNST+NAc KD mice did not differ from controls.

**Figure 5:**
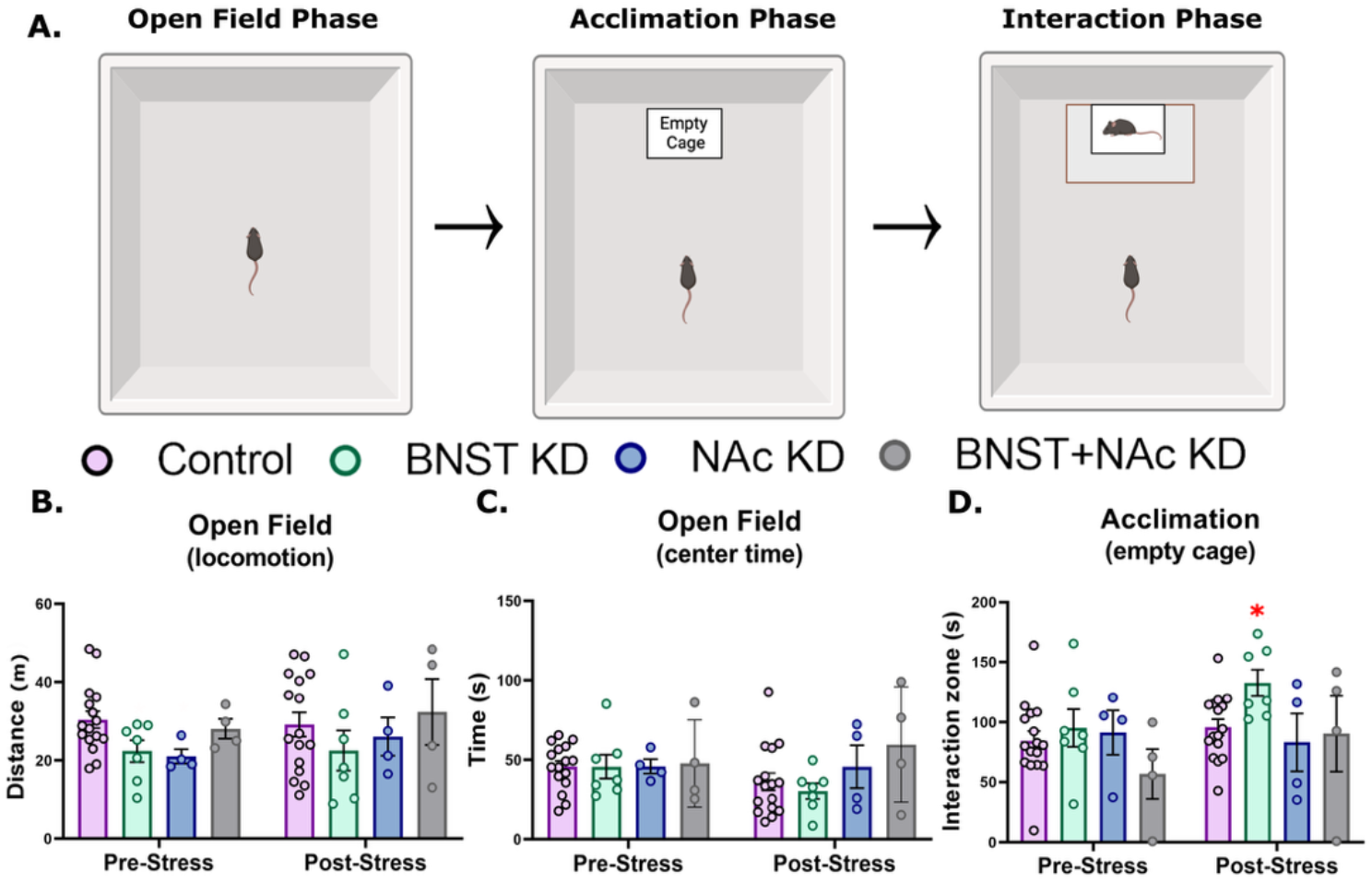
Large arena test open field and acclimation phase data. (A) Schematic illustration of the large arena test and the 3 phases: Open Field, Acclimation, and Interaction. (B) Open field locomotion activity for all treatment groups before and after stress (C) Open field center time plotted for all treatment groups before and after stress (D) Acclimation interaction time with an empty cage for all treatment groups before and after stress* P < 0.05 vs control.

### OTR Knockdown Does Not Blunt Effects of Stress in the Small Arena

Across all behaviors we quantified, there were no effects of OTR knockdown after stress compared to controls. The only effects of OTR knockdown were observed pre-stress in affiliative nose-to-nose sniffing with naïve target mice (Fig. 6H, X^2^(3)=17.84, p=0.0005), with reduced sniffing in BNST (p= 0.0007, d=1.48) and BNST+NAc KD (p=0.007, d=1.21) compared to controls before stress. There were no effects of OTR knockdown on more assertive anogenital sniffing data before stress. However anogenital sniffing was significantly reduced post-stress in controls with the naïve target mice (Fig. 6D, p=0.0002, d=1.14) but not in other groups. A similar effect of stress on anogenital sniffing was observed post-stress with aggressive target mice (Fig 6I, p=0.02, d=0.70).

**Figure 6:**
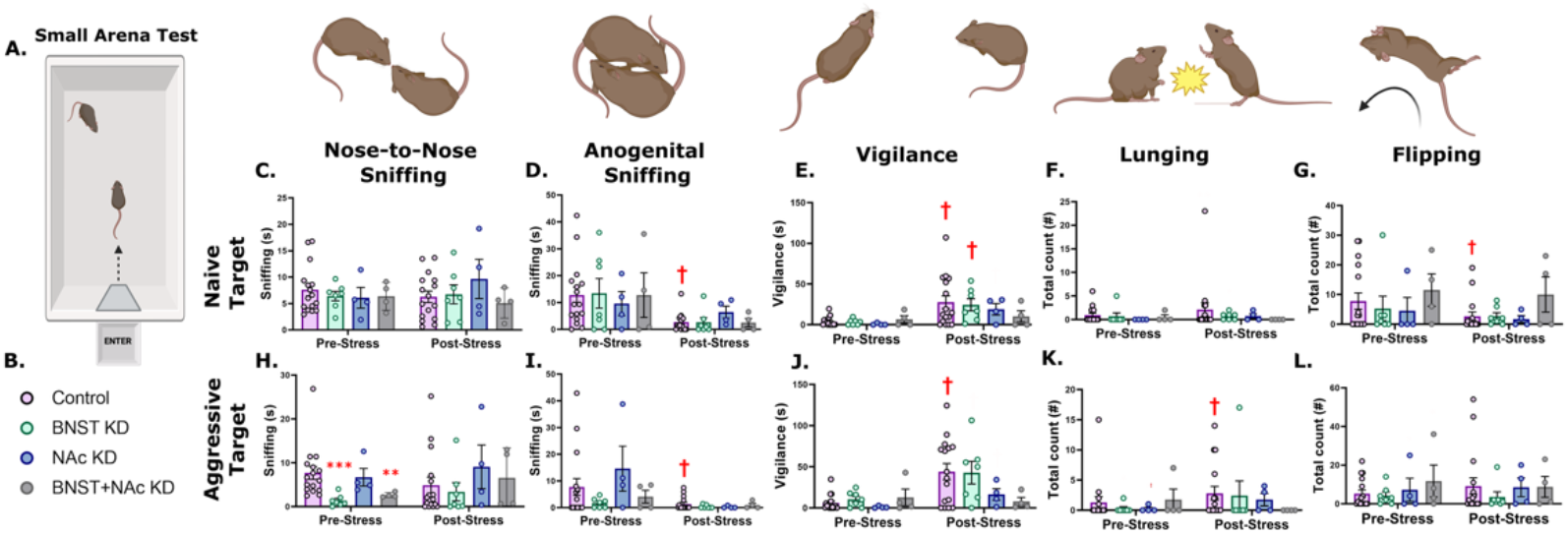
Small arena test social behaviors before and after stress. (A) Schematic illustration for the small arena test. Legend for treatment groups. Nose-to-nose sniffing (C), anogenital sniffing (D), social vigilance (E), lunging (F), and flipping (G) were scored with non-aggressive naïve target mice and then aggressive target mice. Tests were conducted once before social defeat and once after social defeat. † vs. pre-stress paired comparison, ** P < 0.01 vs. control, *** P < 0.001 vs. control.

There were also no treatment effects on vigilance before or after stress (Fig. 6E and J, Target: X^2^(3)=1.92, p=0.59; VP: X^2^(3)=3.07, p=0.38). After stress in paired comparisons, vigilance directed towards naïve target mice was increased in control (p=0.007, d=1.07) and BNST KD (p=0.0156, d=1.51) mice compared to pre-stress observations (Fig. 6E). Similar results were observed with vigilance directed towards aggressive target mice, as stress increased vigilance in control mice (p=0.005, d=1.33) but not in the BNST KD group (p=0.09). No differences were observed in lunging behaviors (Fig. 6F andK). There were no effects of OTR knockdown on flipping behavior (Fig. 6G and L) although reduced flipping was observed post-stress in control animals interacting with naïve target mice (Fig. 6G, p=0.032, d=0.59).

## DISCUSSION

Using CRISPR/Cas9 we observed that experimentally reduced OTR expression in the BNST of female California mice blocked stress-induced decreases in social approach. Additionally, BNST knockdown of OTRs increased approach to a novel object after stress. Unexpectedly, effects of BNST OTR knockdown had more modest effects on social vigilance in the large arena and small arena. These observations contrast with pharmacological studies where BNST OTR antagonists increased social approach and decreased social vigilance in stressed female California mice (Duque-Wilckens et al., 2018) and infusions of OTR agonists decreased approach and increased vigilance in unstressed males and females (Duque-Wilckens et al. 2020; Luo et al., 2022). In contrast knockdown of OTR in NAc significantly reduced stress-induced social vigilance without affecting social approach in the large arena. Previous pharmacological studies reported that activation of OTR in NAc in stressed females increased social approach and decreased vigilance (Williams et al., 2020). A key difference from our previous pharmacology studies is that here OTR manipulations were performed before stress exposure. This raises the possibility that activation of OTR during stress exposure contributes to behavioral changes in female California mice. Overall, our findings suggest that local BNST OTRs modulate stress-induced decreases in social approach.

### Effects of oxytocin receptor on social approach

Our analyses of viral spread and autoradiography indicate that mice assigned to the BNST knockdown had reduced OTR expression selectively in the BNST. This suggests that behavioral effects in this group are mediated by locally expressed receptor on post-synaptic neurons or glia. This finding aligns with prior pharmacological studies where social approach is reduced in stress naïve female California mice when oxytocin receptor agonists are infused into the BNST (Duque-Wilckens et al., 2020; Luo et al., 2022), and activation of OTR in the BNST after stress decreases approach (Duque-Wilckens et al., 2018). Notably, BNST OTR knockdown also increased approach to an empty cage after stress, suggesting that OTR in BNST can modulate exploratory behavior in non-social contexts. In California mice, social defeat generally does not affect approach to the empty cage in the acclimation phase (Trainor et al. 2011 and 2013; Greenberg et al. 2014; Duque-Wilckens et al. 2018). However, males and females treated with an infusion of 1 ng of oxytocin into the BNST showed avoidance of the empty cage (Duque-Wilckens & Trainor unpublished data). It is possible that defeat may not influence approach to an empty cage because oxytocin is not released in this context. In the small arena the effects of OTR knockdown in BNST were more limited when focal mice could freely interact with target mice. Knockdown of OTR in the BNST reduced nose-to-nose sniffing of aggressive target mice before stress, but not after stress. Thus, OTR in BNST may play a more important role in modulating social interactions when individuals are not in close contact. It’s possible that arginine vasopressin (AVP) may play a stronger role in modulating behavior when individuals are in close contact. Vasopressin can activate V1a receptors and previous studies reporting effects of V1 receptor regulation of behavior involve animals physically interacting with one another such as pair bonding (Winslow et al., 1993), social communication (Febo & Ferris, 2016; Freeman et al., 2018) and aggression (Duque-Wilckens et al. 2016, Taylor et al., 2022; Terranova et al., 2016).

In the NAc, previous studies in California mice indicated the importance of OTR in regulating social approach. Social defeat stress decreases OTR binding (Duque-Wilckens et al., 2018) and *Oxtr* mRNA (Williams et al., 2020) in the NAc of male and female California mice while OTR antagonists infused into the NAc reduced social approach (Williams et al., 2020). Here knockdown of OTR in the NAc did not reduce social approach before or after stress. Although this comparison is under-powered relative to the BNST group, this observation is consistent with previous studies in male mice documenting a role for presynaptic OTR in the NAc (Dolen et al. 2013; Nardou et al. 2019). Dopaminergic inputs from the ventral tegmental area (VTA) regulate social approach in C57BL/6J mice by encoding reward value and reinforcing social interactions.

Optogenetic activation of VTA dopamine neurons enhances social preference, while inhibition reduces it (Paquelet et al., 2022). Similarly, serotonergic inputs from the dorsal raphe nucleus (DRN) contribute to learning-related behaviors, with DRN → VTA 5H-T neurons preferentially encoding reward-related cues and dynamically regulating social approach based on prior experience. In juvenile C57BL/6J mice, Dolen and colleagues showed that the behavioral effects of OTR antagonists infused into the NAc were mediated by OTR expressed on presynaptic terminals originating from the DRN that directly affect conditioned place preference learning (Dolen et al., 2013). These findings suggest that social approach behaviors in the NAc may be shaped by coordinated inputs from DRN → VTA serotonin neurons and/or VTA dopamine neurons.

### Effects of oxytocin receptor on social vigilance

Effects of OTR knockdown on social vigilance were distinct from effects on social approach. In the large arena, BNST OTR knockdown produced a non-significant trend for reduced vigilance after stress compared to controls. In previous pharmacology studies (which target pre-and post-synaptic receptors), activation of BNST OTR increased social vigilance in both male and female California mice (Duque-Wilckens et al., 2018 and 2020; Luo et al., 2022). Since post-stress social approach was affected by BNST OTR knockdown, we assessed the relationship between social approach and vigilance. If pre-synaptic OTR in the BNST were controlling social vigilance, we would expect that the negative relationship between social approach and vigilance would be eliminated by OTR knockdown (which should not impact pre-synaptic receptors). Approach and vigilance were inversely related in both control and BNST knockdown mice, which does not provide strong evidence for a role of presynaptic OTR in controlling vigilance. We also examined whether the extent of knockdown was correlated with vigilance and observed little evidence that mice with higher vigilance in the BNST KD group had more receptor density (e.g. less effective knockdown). One possibility is that activation of OTR in BNST during episodes of social defeat may help encode the salience of an aversive interaction. Our knockdown approach reduced receptor expression before episodes of social defeat whereas our previous antagonist experiment only inactivated OTR after defeat (Duque-Wilckens et al., 2018). Encoding effects of OTR during episodes of social stress may be stronger in the NAc.

Knockdown of OTR in the NAc significantly reduced vigilance after stress in the large arena, even though there was less statistical power due to smaller sample size than the BNST group. Similar patterns were observed in the small arena. Suppressed vigilance in the NAc knockdown group was an unexpected finding but could reflect a learning component. In the NAc, interactions between oxytocin and dopamine are known to modulate social behaviors (Liu & Wang, 2003). There is strong evidence that dopamine signaling in the NAc organizes responses to aversive experiences, with D2 receptor-expressing medium spiny neurons in the core supporting the encoding and recall of aversive foot-shocks (Belilos et al., 2023). In Syrian hamsters, dopamine release in the NAc core increases during both winning and losing aggressive encounters (Cross et al., 2025), and pharmacological blockade of dopamine receptors in the NAc during losing blocks increases in submissive behavior in future social encounters (Gray et al., 2015). This suggests that effects of OTR knockdown in NAc on vigilance behavior, similar to dopamine receptor signaling, may depend on its timing relative to social defeat. Overall, these findings raise the possibility that the NAc is a site where oxytocin and dopamine interact during stress to influence vigilance behaviors.

## CONCLUSIONS

These results demonstrate that locally expressed OTRs in the BNST and NAc make distinct contributions to stress-induced social behaviors. In the BNST, OTR knockdown decreased social avoidance and increased exploratory behaviors after stress, while effects on vigilance were modest. Effects of OTR knockdown in the NAc on vigilance suggest a possible role for oxytocin in the mesolimbic dopamine system for encoding socially aversive experiences. Together, these findings add to evidence from pharmacology studies that suggest that BNST and NAc OTRs have distinct effects on social approach and vigilance. Further work is needed to determine how these populations of OTR act during episodes of stress versus modulating the long-lasting effects of stress on behavior.

## ACKNOWLEDGEMENTS

The authors thank H. Butler-Struben, PhD and Pei Luo, PhD for their help in surgical training and optimizing the micro-injection protocol for this study. Illustrations for figures were done with biorender.com. This work was supported by an Ikerbasque Research Fellowship to AJB, P50MH100023 and R01MH112788 to LJY, OD P51OD011132 to EPC (Primate center base grant), and NIMH R01MH121829 and R01MH121829-04S1 to BCT.

